# Thetaburst TMS to the posterior superior temporal sulcus decreases resting-state fMRI connectivity across the face processing network

**DOI:** 10.1101/794578

**Authors:** Daniel A Handwerker, Geena Ianni, Benjamin Gutierrez, Vinai Roopchansingh, Javier Gonzalez-Castillo, Gang Chen, Peter A Bandettini, Leslie G Ungerleider, David Pitcher

## Abstract

Humans process faces using a network of face-selective regions distributed across the brain. Neuropsychological patient studies demonstrate that focal damage to nodes in this network can impair face recognition, but such patients are rare. We approximated the effects of damage to the face network in neurologically normal human participants using thetaburst transcranial magnetic stimulation (TBS). Multi-echo functional magnetic resonance imaging (fMRI) resting-state data were collected pre- and post-TBS delivery over the face-selective right superior temporal sulcus (rpSTS), or a control site in the right motor cortex. Results showed that TBS delivered over the rpSTS reduced resting-state connectivity across the extended face-processing network. This connectivity reduction was observed not only between the rpSTS and other face-selective areas, but also between non-stimulated face-selective areas across the ventral, medial and lateral brain surfaces (e.g. between the right amygdala and bilateral fusiform face areas and occipital face areas). TBS delivered over the motor cortex did not produce significant changes in resting-state connectivity across the face-processing network. These results demonstrate that, even without task-induced fMRI signal changes, disrupting a single node in a brain network can decrease the functional connectivity between nodes in that network that have not been directly stimulated.

**Author Summary:** Human behavior is dependent on brain networks that perform different cognitive functions. We combined thetaburst transcranial magnetic stimulation (TBS) with resting-state fMRI to study the face processing network. Disruption of the face-selective right posterior superior temporal sulcus (rpSTS) reduced fMRI connectivity across the face network. This impairment in connectivity was observed not only between the rpSTS and other face-selective areas, but also between non-stimulated face-selective areas on the ventral and medial brain surfaces (e.g. between the right amygdala and bilateral fusiform face areas and occipital face areas). Thus, combined TBS/fMRI can be used to approximate and measure the effects of focal brain damage on brain networks, and suggests such an approach may be useful for mapping intrinsic network organization.

**Technical Terms:** *TBS vs TMS:* Transcranial magnetic stimulation (TMS) is a method that induces current in neural tissue by using a rapidly changing magnetic field. The pattern of magnetic field changes can vary. Thetaburst TMS (TBS) is a type of TMS where the same stimulation pattern fluctuates at around a 5Hz cycle.

*Multi-echo fMRI:* During typical fMRI, protons are excited and there is a delay, the echo time, before data are collected. That delay is typically designed to result in a high contrast for blood oxygenation differences. In multi-echo fMRI, data are collected at several echo times each time protons are excited. This results in data that have different levels of contrast for blood oxygenation differences. This added information can be used to empirically decrease noise.

*Face network:* A group of brain regions that show significant activity changes in response to visual face stimuli. While these regions have been defined using univariate analyses with task-based fMRI, they often significantly correlate with each other at rest. In this manuscript, the following regions were a priori defined as part of the face network: posterior superior temporal sulcus (pSTS), amygdala, fusiform face area (FFA), and occipital face area (OFA).

*Matrix based analysis (MBA):* A recent approach that uses a Bayesian multilevel modeling framework to identify pairs of ROIs where a decrease in correlation magnitude was larger than expected along with a measure of statistical evidence. With this approach, correlations between all pairs of ROIs are assessed as part of a single model rather than many independent statistical tests.

## Introduction

The ubiquitous presence of faces in our environment makes them a salient stimulus for studying the cognitive functions of the human brain. Functional magnetic resonance imaging studies have identified regions across the brain that exhibit a stronger neural response to faces than to objects (Gauthier et al., 2000; Kanwisher, McDermott, & Chun, 1997; McCarthy, Puce, Gore, & Allison, 1997; Phillips et al., 1997; Puce, Allison, Bentin, Gore, & McCarthy, 1998). These face-selective regions are linked to form distributed nodes of a face-processing network (Calder & Young, 2005; Haxby, Hoffman, & Gobbini, 2000). Neuropsychological patients with damage to these face-selective areas exhibit face-selective recognition impairments (Barton, 2008; Bouvier & Engel, 2006; Landis, Cummings, Christen, Bogen, & Imhof, 1986; Rezlescu, Barton, Pitcher, & Duchaine, 2014; Rossion et al., 2003), providing strong evidence that faces are processed in a specialized network. Despite the importance they have afforded the study of brain function, such patients are rare. In addition, the interpretation of the data they produce is limited by individual differences in pre-morbid ability (Farah, 2004), and compensatory effects of plasticity that may have occurred after the incident (Robertson & Murre, 1999).

In the present study we used thetaburst transcranial magnetic stimulation (TBS) in combination with multi-echo fMRI acquisition (Posse et al., 1999). Resting-state fMRI was used as a proxy for the impact of disruption on one node in the face network of neurologically normal experimental participants. Over two sessions, these participants were scanned using multi-echo fMRI pre- and post-TBS delivery over the face-selective right superior temporal sulcus (rpSTS), the stimulation site of interest, or the hand area of the right motor cortex, the control site. Multi-echo fMRI denoising is a relatively new method that quantitatively identifies and removes non-blood oxygen-weighted noise from fMRI data (Kundu et al., 2013). Prior to TBS, participants viewed 3-second videos of moving faces and objects (Pitcher, Dilks, Saxe, Triantafyllou, & Kanwisher, 2011) during fMRI scanning to functionally localize face-selective regions of interest (ROIs).

The face-selective areas in the rpSTS, the fusiform face area (FFA) (Kanwisher et al., 1997; McCarthy et al., 1997), and the occipital face area (OFA) (Gauthier et al., 2000) comprise the core nodes of the face processing network (Calder & Young, 2005; Haxby et al., 2000). In addition to these core nodes, fMRI studies have also identified face-selective voxels in the amygdala (Phillips et al., 1997). Face processing models propose these areas perform different cognitive functions (e.g. recognizing identity, facial expression or eye gaze direction), but share task-relevant information. The existence of functional connections between face areas is supported by evidence showing that disruption to one of these areas can impair a range of face recognition tasks. For example, TMS delivered over the rpSTS disrupts the McGurk effect (Beauchamp, Nath, & Pasalar, 2010) as well as performance on face-processing tasks involving eye-gaze (Pourtois et al., 2004), trustworthiness judgments (Dzhelyova, Ellison, & Atkinson, 2011) and discriminating facial expressions (Pitcher, 2014; Sliwinska & Pitcher, 2018). Neuropsychological patients with lesions to the rpSTS also show face processing impairments. A patient with a lesion to the rpSTS and angular gyrus was impaired on an unfamiliar face matching task (Sakurai, Hamada, Tsugawa, & Sugimoto, 2016), while another patient with a lesion to right superior temporal gyrus was impaired at discriminating gaze detection (Akiyama et al., 2006).

Previous studies have shown TMS-induced face network activation changes during tasks (Pitcher, Duchaine, & Walsh, 2014; Pitcher, Japee, Rauth, & Ungerleider, 2017). The nodes of the face network also show functional integration during resting state fMRI (Li, Song, & Liu, 2019; X. Wang et al., 2016; Zhang, Tian, Liu, Li, & Lee, 2009). In this study, we tested whether the same face network areas defined in those studies show correlation decreases in response to TMS to the rpSTS, even in the absence of a face stimulus. If so, then this would build evidence that behaviorally defined face processing regions are part of a network whose nodes regularly communicate and interact with each other without this communication being induced by a specific set of stimuli. Results demonstrated that TBS delivered over the rpSTS caused a reduction in correlations between the stimulated node and unstimulated nodes of the face processing network.

## Results

For each volunteer, face-selective regions of interest (ROIs) were localized using voxels with larger responses to face videos than to object videos for the left posterior superior temporal sulcus (lpSTS), bilateral fusiform face areas (FFA), bilateral occipital face areas (OFA) and bilateral amygdala. The right posterior superior temporal sulcus (rpSTS) ROIs were face-selective voxels in gray matter that were also within an 18mm diameter sphere centered on the stimulation site. This added restriction means the voxels in the rpSTS ROI were likely to have been directly affected by TBS stimulation. The stimulation site was defined using the same functional localizer scan, but collected on a preceding session. The right hand motor stimulation site was anatomically defined using an anatomical scan from a preceding session. ROIs for the bilateral hand areas were manually drawn following the gray matter anatomy of the hand knob in the precentral sulcus. The locations for the regions of interest are shown in Figure S1.

For ROI-based analyses, mean time series of the ten-minute resting state fMRI data were calculated using the voxels within each ROI. The correlations between these ROIs were calculated for the resting runs pre- and post-TBS stimulation. The correlation coefficients were Fisher Z transformed. Since the ROIs are *apriori* selected from a network where we expect to see disruption (Pitcher et al., 2014; Pitcher et al., 2017) – with the hand motor regions as a control, the statistical changes of interest will focus on the ROI analyses. A matrix based analysis (MBA) through Bayesian multilevel modeling was used to identify pairs of ROIs where a decrease in correlation magnitude was larger than expected along with a measure of statistical evidence (Chen et al., 2019). The advantage of this approach is that, instead of adopting a univariate GLM with the assumption that each ROI pair is an independent entity that shares no commonality or similarity with its peers, the magnitude estimates and uncertainties of all ROI pairs are assessed as part of a single integrative model.

Figure 1 shows the effects of stimulation on the correlations pre – post TBS for all pairs of ROIs (bilateral STS, FFA, OFA, amygdala and hand motor cortices) using Bayesian multilevel modeling. For rpSTS stimulation, multiple ROI pairs within the face processing network showed decreased connectivity that would be likely to be greater than zero with a posterior probability of at least 0.95 (Fig 1A). For pre – post TBS to the motor cortex (Fig 1B) and interaction effects, there was no strong evidence that any ROI pairs were likely to be greater than zero. All posterior probabilities were less than 0.85 and all effect sizes were less than 0.07 (Fig. S2).

**Figure 1:**
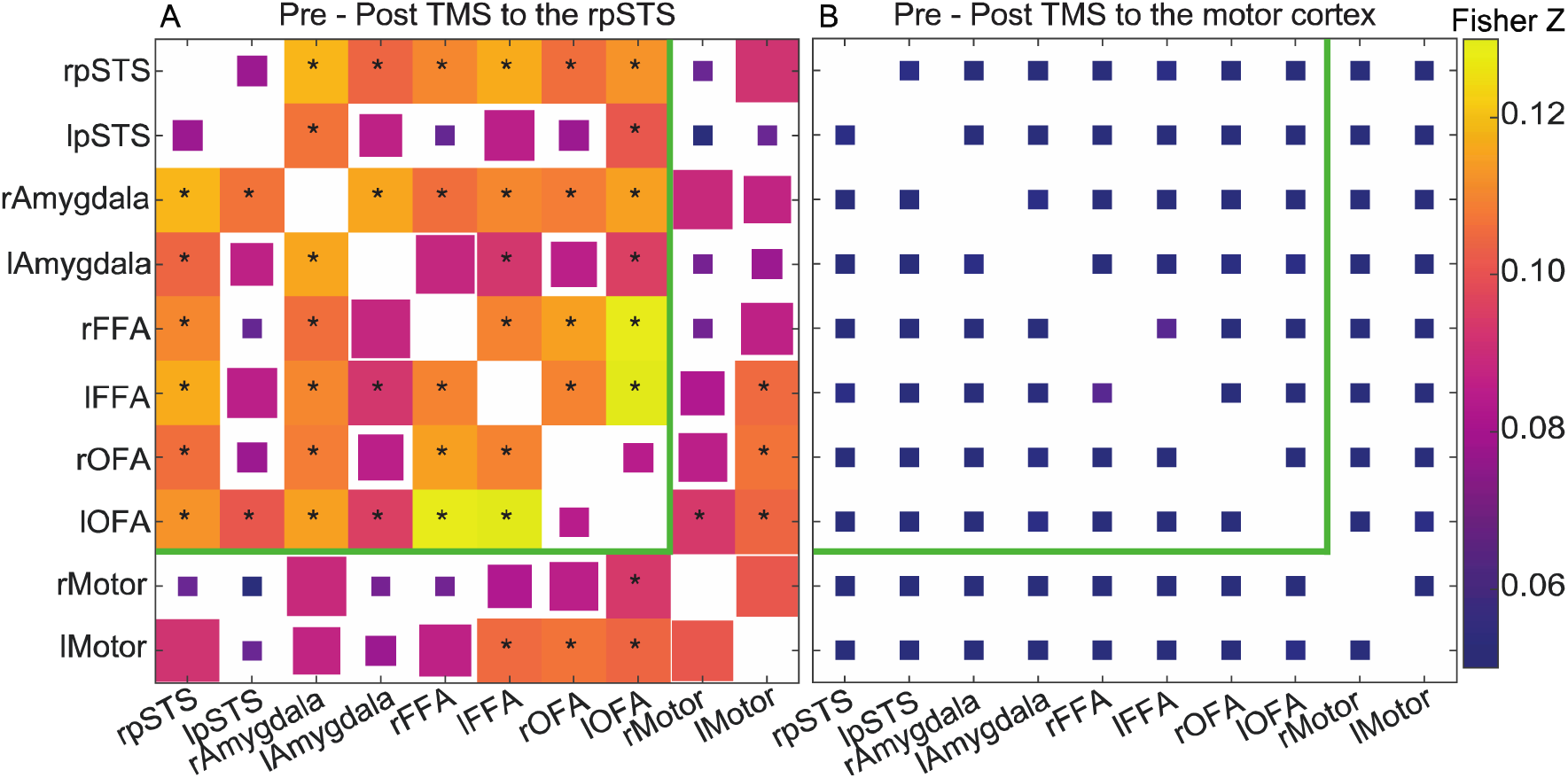
Fisher Z transformed correlation magnitude changes from pre - post TMS to the (A) rpSTS and (B) right motor cortex. Magnitudes are the Matrix Based Analysis (MBA) model fits across the population. Changes that are likely to be greater than 0 with a posterior probability of at least 95% are full squares and include an *. The squares’ edge lengths decrease linearly to their minimum size for a 90% likelihood or less. The green line marks the ROI pairs that are within the pre-defined face-selective network. Fig. S2 contains an unthresholded version of this figure and the interaction effect.

To help visually compare the effect sizes for rpSTS vs motor stimulation, Figure 2 shows boxplots for the correlations between the rpSTS and the other ROIs for the runs pre and post TBS. The colored section of the pie charts in this figure show the number of volunteers that had a pre – post TBS correlation decrease. These pie charts show that 11-13 of the 16 volunteers showed a correlation decrease post rpSTS stimulation. The supplementary material includes boxplots for correlation magnitudes of all ROI pairs (Figs S3 and S4). Whether the correlation is between a pair of ROIs that includes the directly stimulated rpSTS or another pair of face network ROIs, the correlations to other ROIs in the face network decreased across the majority of volunteers when the rpSTS was stimulated.

**Figure 2:**
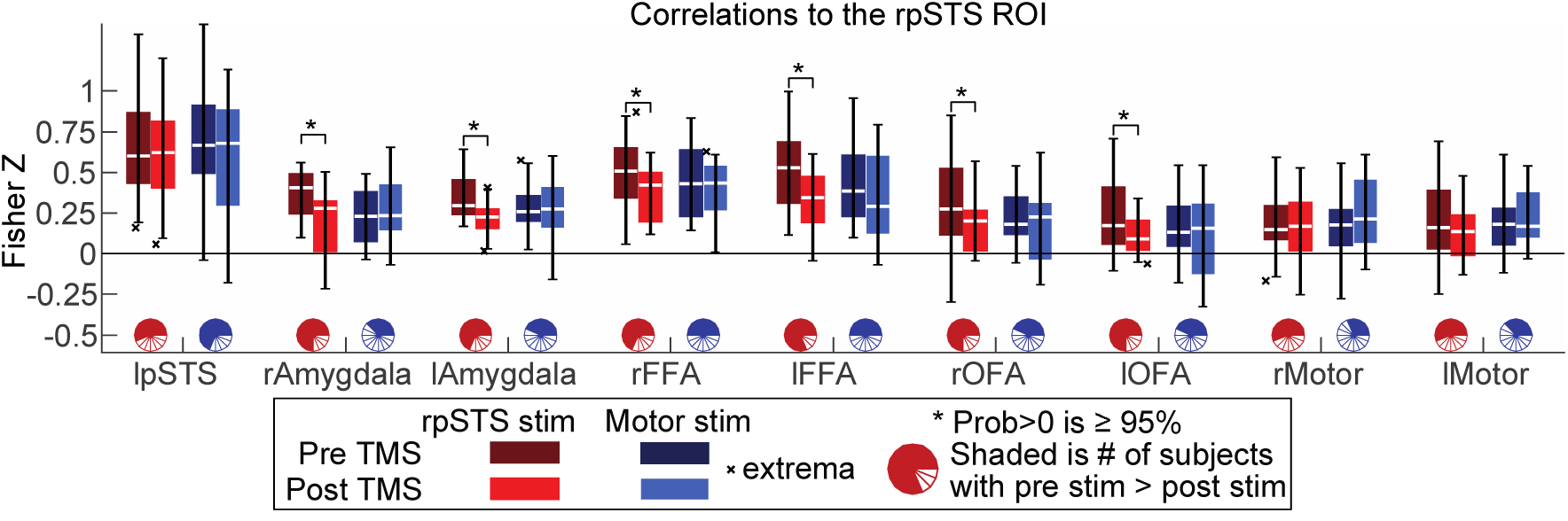
Correlations between the rpSTS ROI and ROIs in a pre-defined face network as well as the bilateral primary motor hand regions. These data correspond to the first row of the matrix in figure 1. The full matrix of bar plots in in Fig. S2. Magnitudes are the MBA model fits across the population for each condition. Boxplots show 25-75% of the distribution. The white line is the median. Whiskers are the maximum and minimum values excluding outliers. MBA used to calculate posterior probabilities that a difference is greater than 0. The correlations between the rpSTS and other face regions consistently decrease post rpSTS stimulation, but not post rMotor stimulation.

There is a possibility that TBS to the rpSTS would cause correlation decreases between the rpSTS and the entire brain. To test for this possibility, all subjects’ data were aligned to each other and a group map was calculated with an ANOVA using multivariate modeling (Figure 3). While there are decreases in activity that do not all cross meaningful statistical thresholds both within and outside the face network, the correlation decreases post rpSTS stimulation are clearly not a global effect.

**Figure 3:**
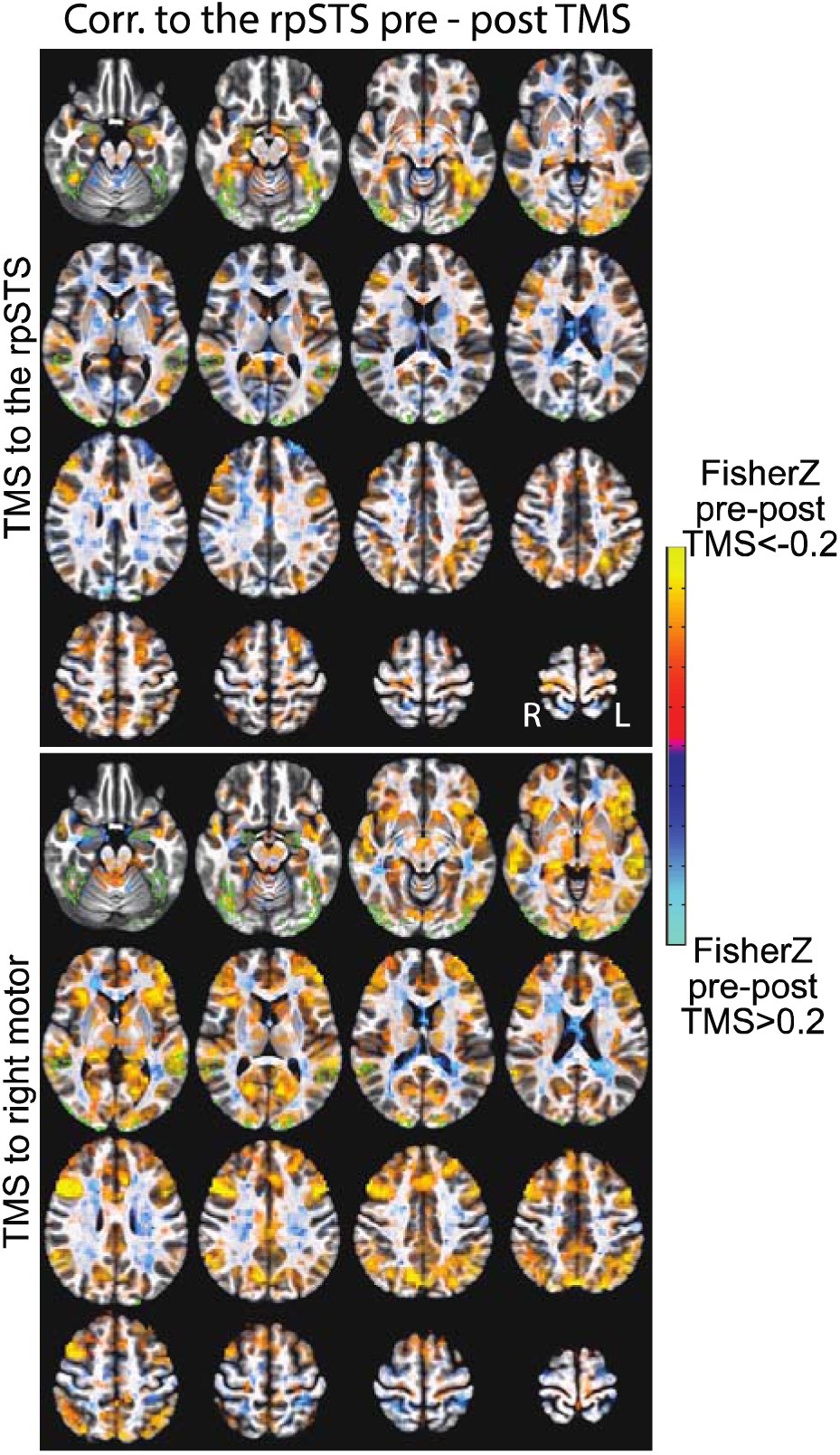
Whole brain correlation changes (Fisher Z transformed) pre - post TBS stimulation. Voxel coloring shows the magnitude of the correlation difference. Voxels are opaque for p>=0.01 to transparent for p=0.The green outlines are voxels in the ROIs of the pre-defined face network in at least 3 volunteers. For TMS to the rpSTS, many of the larger signal drops are within the face network ROIs. TMS to right motor shows decreases across more of the brain that isn’t localized to the face network.

## Discussion

Models of the human face processing network propose that face-selective areas perform different functional roles but share task-relevant information when processing faces (Calder & Young, 2005; Haxby et al., 2000). In the current study, we approximated the impact of disruption to the face network by delivering TBS over the face-selective area in the right posterior superior temporal sulcus (rpSTS). The impact of the TBS stimulation was measured with resting-state fMRI, with the goal of examining whether the relationship between face network regions is altered even in the absence of a face-selective cognitive task. Our results demonstrated that disruption of the rpSTS reduced resting-state connectivity, measured by correlated activity, across the nodes of the face processing network. Figure 1 shows a reduction in connectivity between the rpSTS and bilateral FFA, OFA, and amygdala as well as between 14 of 21 pairs of ROIs in the face network that were not directly stimulated. This result demonstrates that disrupting a single face-selective area causes widespread connectivity decreases across the extended face processing network.

In addition, functional connectivity between different nodes of the face network has been shown to correlate with behavioural measures of face recognition. Zhu and colleagues demonstrated that connectivity between the OFA and FFA correlated with performance on face, but not object, matching discrimination tasks (Zhu, Zhang, Luo, Dilks, & Liu, 2011). O’Neill and colleagues demonstrated that connectivity between the FFA and perirhinal cortex correlated with an upright, but not inverted, face matching task (O’Neil, Hutchison, McLean, & Kohler, 2014). Most recently, Ramot and colleagues demonstrated that face recognition memory was correlated with connectivity between the face network and regions of the medial temporal lobe (including the hippocampus) (Ramot, Walsh, & Martin, 2019). Taken together, these findings suggest that synchronized spontaneous neural activity between face regions might be responsible for the different behavioural aspects of face recognition. The current study builds upon and extends these prior findings to show that transient disruption of a single node in the face network can disrupt functional connectivity across the wider network.

Figures 2, S3, and S4 visualize the effect of TBS across subjects to show that correlations between face network regions have correlation decreases for the majority of volunteers post rpSTS stimulation. This widespread pattern of distributed correlation decreases suggests that the nodes of the face network are tightly interconnected, and that distruption of one node can affect other nodes in the network. Importantly, TBS delivered over the right motor cortex (which acted as a control site) did not produce above chance reductions in functional connectivity across the face network. The observed network-wide drop in resting-state connectivity is consistent with models proposing that the rpSTS is a core component of the face-processing network (Calder & Young, 2005; Haxby et al., 2000). While we do show an effect of rpSTS stimulation and not an effect for motor stimulation, we do not observe a significant effect of motor cortex stimulation between motor ROIs, nor an interaction effect. Our results do not completely match our predictions, possibly due to the combined effect of two inherently noisy techniques, namely, resting-state fMRI and TBS. Resting-state fMRI is known to be more strongly affected than task fMRI by many noise sources (e.g. inter-subject cognitive variation, physiological noise, head motion, scanner instabilities). Also, there is still not a full understanding of all factors that modulate TBS effect size including how much spatial precision matters, whether different brain regions are more sensitive to TBS than others, and the causes of inter-subject variability in TBS sensitivity. Specifically in this study, we note our post-stimulation observation of spatial variation between the hand motor ROI and the hand motor stimulation site (Fig S1). This variation in stimulation site may have led to an observed increase in variance of correlation changes post motor stimulation vs rpSTS stimulation (Fig S3 and S4). That added variance may have reduced the posterior probabilities involving motor stimulation data. Although we did our best to account for known sources of noise, residual artifacts can be expected to exist, and those may have a multiplicative effect when combining the techniques, further hindering our statistical power.

Additionally, figure 3 shows large, but not statistically significant, correlation decreases in response to TBS to the rpSTS. These occur both within and outside the a priori defined face network regions. While we will not attempt to interpret these below-threshold correlation decreases, it is important to note that rpSTS function is not limited to face processing. Additionally, the stimulation of an approximately 1cm^3^ volume of cortex may affect brain regions outside of a face network.

In this study, we could not directly test whether TBS to the rpSTS caused alterations in face processing because we cannot both collect resting state fMRI and also present face-selective stimuli during the post TBS time window when the TBS effect is observable. However, in previous studies using face-selective stimulus evoked BOLD mmeasures and the same TBS stimulation protocol as used here, changes were reported in responses to face stimuli (Pitcher et al., 2014; Pitcher et al., 2017). The reduction in network connectivity we observed between non-stimulated face-selective regions on the medial and ventral brain surfaces extends the findings of this prior work of Pitcher et al. Our previous combined TBS/fMRI studies examined the impact of disruption in face-selective regions while participants viewed face videos. These studies showed that TBS delivered over the rpSTS reduced the neural response to faces in the rpSTS itself, as well as in other face-selective areas including the right fusiform face area (rFFA) (Pitcher, 2014) and face-selective voxels in the right amygdala (Pitcher et al., 2017). However, while these studies reported a reduction in the neural response to faces in remote face-selective areas, neither study was capable of detecting changes in functional connectivity between face-selective areas. While prior studies demonstrated functional connectivity between the FFA and the amygdala (Fairhall & Ishai, 2007; Herrington, Taylor, Grupe, Curby, & Schultz, 2011; Vuilleumier, Armony, Driver, & Dolan, 2001, 2003), our study shows that correlations between these areas are reduced post-TBS delivery over a remote node in the network, in this case the rpSTS.

Non-human primate neuroanatomical studies in macaques report a cortical pathway projecting from the STS to dorsal sub-regions of the amygdala (Aggleton, Burton, & Passingham, 1980; Stefanacci & Amaral, 2000, 2002). In addition, neuroimaging and lesion studies of humans and macaques demonstrate that the amygdala is engaged in facial expression recognition (Adolphs, Tranel, Damasio, & Damasio, 1994; Adolphs et al., 1999; Calder et al., 1996; Hadj-Bouziane et al., 2012; Hoffman, Gothard, Schmid, & Logothetis, 2007; Morris et al., 1996). We previously proposed that humans have a cortical pathway projecting along the STS into the amygdala that processes changeable facial aspects (e.g expression), which was supported using combined TBS/fMRI during face, body, and object viewing (Pitcher et al., 2017). By replicating this effect using functional connectivity and without a viewing task, we provide further support for a pathway between these regions.

Neuropsychological patients exhibiting face-selective recognition impairments were essential to the development of cognitive and brain models of face processing (Bruce & Young, 1986; Haxby et al., 2000). Studies of patients with right lateral lesions (in the area of the pSTS) showing impairments with eye gaze direction detection (Akiyama et al., 2006; Campbell, Heywood, Cowey, Regard, & Landis, 1990) and unfamiliar face identity matching (Sakurai et al., 2016) have been reported. However, prosopagnosic patients with lateral lesions are less common than those with ventral lesions. Our study further demonstrates that transiently disrupting the brains of neurologically normal experimental participants with TBS offers an alternative and safe proxy for modeling the effects of focal cortical disruption on cognitive networks. TBS can be combined with functional magnetic resonance imaging (fMRI) to study the networks that process scenes (Mullin & Steeves, 2013), faces (Rafique, Solomon-Harris, & Steeves, 2015; Solomon-Harris, Rafique, & Steeves, 2016), decision making (Rahnev, Nee, Riddle, Larson, & D’Esposito, 2016) and memory (J. X. Wang et al., 2014). Crucially, while we do see some decreases between a few face network ROIs and the motor ROIs (Fig 1), we examined the voxel-wise correlations across the brain to show that the largest distruptions of TBS to the rpSTS were in the face selective network. Still, the decreases in some connections to motor cortex ROIs hint at how altering a single node can potentially distrupt multiple distinct networks. Each brain region does not contribute to only one self-contained network of brain regions.

Neurologists have studied the impact of focal brain lesions in patients for over two hundred years. The study of these patients has been highly influential and has produced insights into human cognition. However, patients with focal lesions in brain regions of interest are extremely rare. In addition, the interpretation of the data they produce is tempered by factors like individual differences in pre-morbid ability (Robertson & Murre, 1999) and the unknown effects of neural plasticity that may have occurred after their incident (Farah, 2004). TBS combined with resting state fMRI has been used to study the default mode network (Abellaneda-Perez et al., 2019; Shang et al., 2019), attention (Anderkova et al., 2018), visualspatial neglect (Fu et al., 2017), cerebellar connectivity (Rastogi et al., 2017), mental illness, (Baeken, Duprat, Wu, De Raedt, & van Heeringen, 2017) and memory in humans (Mancini et al., 2017). These studies show that measuring transient TBS induced disruption with resting state fMRI offers a systematic experimental methodology for extending more than two hundred years of neuropsychological research.

In conclusion, models propose that face-selective brain areas can be linked together to form the distributed components of a face processing network (Calder & Young, 2005; Haxby et al., 2000). The current findings support these models by showing that transient disruption to one of these components, the rpSTS, results in widespread correlation decreases across the distributed nodes of this network that persists for ten minutes.

## Materials and Methods

### Participants

17 right-handed participants (11 female, age mean±stdev=27±5), with no major medical illness, no neurological or psychiatric illness, no serious head injury, no learning disability, no history of drug or alcohol abuse in the past 3 months, no prescription drugs or supplements affecting brain function, no serious vision or hearing problems and with normal, or corrected-to-normal, vision gave informed consent as directed by the National Institutes of Health (NIH) Institutional Review Board (ClinicalTrials.gov identifier: NCT01617408). One participant was excluded due to high motion (207 of 1200 volumes in the 4 resting runs censored due to motion or outliers vs 0-40 for all other volunteers).

### Experimental Design

#### Procedure

Participants completed three separate fMRI sessions, each performed on a different day. The first session was an fMRI experiment designed to individually localize the TBS stimulation sites in each participant. Participants viewed face and object videos and a high resolution structural scan was also taken. The data collected in this initial session were used for TBS target site identification only. During the two subsequent fMRI sessions, participants were scanned pre and post receiving TBS stimulation of either the right posterior superior temporal sulcus (rpSTS) or the right primary motor cortex hand knob (rMotor). Stimulation site order was balanced across participants. The two TBS sessions were 7 – 182 days apart (median=23). All the data presented were collected during these two TBS sessions.

#### Stimulation Site Localizer Session and Selection

Stimulation sites were localized using individual structural and functional images collected during an fMRI localizer task that each participant completed prior to the combined TBS/fMRI sessions. For the independent localizer runs used to identify face-selective regions of interest (ROIs), participants viewed 3-second video clips of faces, objects, and scrambled objects. There were sixty movie clips for each category in which distinct exemplars appeared multiple times. Movies of faces were filmed on a black background, and framed close-up to reveal only the faces of 7 children as they danced or played with toys or adults (who were out of frame). Fifteen different moving objects were selected that minimized any suggestion of animacy of the object itself or of a hidden actor pushing the object (these included mobiles, windup toys, toy planes and tractors, balls rolling down sloped inclines). Within each block, stimuli were randomly selected from within the entire set for that stimulus category. This meant that the same actor or object could appear within the same block. These stimuli were used in a previous fMRI study of face perception (Pitcher et al., 2011) and previous combined TBS/fMRI studies (Pitcher et al., 2014; Pitcher et al., 2017).

The rpSTS was identified using a contrast of faces greater than objects. The rpSTS stimulation target was the peak voxel in the significant cluster in the rpSTS. A T1 weighted anatomical scan was also collected during this session and the right motor cortex was the most superior location on the motor cortex hand knob. A posthoc re-examination identified two vounteers where the motor stimulation zone may not have directly stimulated the hand motor region (Fig S1). This may have contributed to a greater variance of responses to hand motor stimulation across the population and thus the lower posterior probabilities for correlation decreases post motor stimulation and interaction effects between the stimulation sites.

#### Combined fMRI/TBS sessions

Data acquisition: Participants were scanned using a GE 3-Tesla MR 750 scanner at the National Institutes of Health (NIH). fMRI images were acquired using a 32-channel head coil (36 slices, 3 × 3 × 3 mm, FOV=21.6 cm, grid = 72×72, flip angle = 77°, ASSET=3, TR = 2 s, TEs = 14.8, 27.1, & 39.5 ms). In addition, high-resolution MPRAGE anatomical scans (T1-weighted, 1 x 1 x 1 mm resolution) were acquired to anatomically localize functional activations and register functional data between sessions.

Before each MRI session, the stimulation site (either rpSTS or rMotor) was located using the Brainsight TMS-MRI coregistration system (Rogue Research), the information was extracted from the subject-specific localizer scans (see the previous “Stimulation Site Localizer Session and Selection” section) and the proper coil locations were marked on each participant’s scalp using a marker pen. During the MRI session, two 10-minute resting-state scans (310 volumes) were acquired followed by functional localizer scans similar to those used for the stimulation site localizer session and then an MPRAGE anatomical scan. The volunteers were then removed from the scanner.

A Magstim Super Rapid Stimulator (Magstim; Whitland, UK) was used to deliver the TBS via a figure-eight coil with a wing diameter of 70 mm. TBS was delivered at an intensity of 80% of active motor threshold or 30% of machine output (whichever was higher) over each participant’s functionally localized rpSTS or the right motor hand knob. We used a continuous TBS paradigm (Huang, Edwards, Rounis, Bhatia, & Rothwell, 2005) of 3 pulses at 50 Hz repeated at 200-ms intervals for a 60-second uninterrupted train of 900 pulses. This is the same protocol that was used in our previous combined TBS/fMRI studies of the rpSTS (Pitcher et al., 2014; Pitcher et al., 2017). We matched the protocol to aid with comparisons across studies. The Stimulator coil handle was held pointing upwards and parallel to the midline.

As soon as stimulation ended, volunteers were returned to the MRI scanner where the following data were acquired: a brief anatomical reference scan to prescribe a slice placement that was visually similar to the pre-TBS scan, an ASSET calibration scan, and two 10-minute resting-state scans. The resting scans began 2.5-5 minutes after the last TBS pulse was delivered depending on how quickly the participant was able to return to the scanner. After the resting scan, another MPRAGE anatomical scan was acquired. Three of the 34 MRI sessions did not include an anatomical scan during the post-TBS part of the session due to time constraints.

The second 10-minute resting scans both pre and post TBS had more head motion in many subjects. For 14 of the 64 pre or post TBS run pairs, there was at least 5 more motion censored volumes in the second run, versus 1 for first run – second run. Similarly the median delta motion (frame-to-frame difference) was larger for 49 of the second runs vs first runs. The average across runs of median delta motion difference between run pairs was 0.01 larger in the second runs. The average median delta motion was 0.05 so this was a 19% increase in median motion. There was only a 7% difference in median delta motion for the first and third runs, which were used in these analyses. While these are subtle differences, the systematic nature of this difference added a confound that made these data difficult to interpret, so the second runs were excluded from all analyses.

## Data processing

Data were processed using AFNI (Cox, 1996). For each fMRI run: The first four pre-steady state volumes were removed; Data were despiked to remove non-biological large signal fluctuations and slice time corrected. Spatial alignment transforms for motion correction within runs and registration across runs were calculated using the middle echo (TE=27.1ms) time series. Non-uniformity of image intensity for the first volume of this time series was corrected using the bias correction code in SPM8. This reference fMRI volume was aligned to the MPRAGE anatomical from the same pre or post TBS session, if available. Otherwise, it was aligned to the anatomical scan from the same day. The anatomical scans were aligned to each other and averaged to make a single, higher quality anatomical volume per volunteer. Registration quality for each pair of anatomical scans and for each fMRI to anatomical alignment were visually inspected for accuracy. The motion correction and alignment transforms were combined into a single affine transform matrix and applied to fMRI time series for all echoes so that fMRI data from each subject was in a single subject-specific space across sessions.

The multi-echo data were then denoised using the MEICA algorithm. MEICA uses the expected properties of signal changes across multiple echoes times to identify and remove signal fluctuations that are unlikely to reflect the blood oxygenation differences that are central to fMRI. By doing this, MEICA has been shown to increase the signal-to-noise ratio and remove data artifacts (Kundu et al., 2013; Kundu, Inati, Evans, Luh, & Bandettini, 2012). The version of the denoising code that we used is available at: https://github.com/handwerkerd/MEICA_FaceNetworkTMS. We will refer to the output of the full MEICA algorithm as “denoised” data. The denoised data will be used for all analyses in this manuscript.

Univariate statistical maps and correlations were calculated within subjects. For functional localizer runs, time series were intensity normalized by dividing by the mean value in each run. For resting-state runs, volumes when the first derivative Euclidian norm of the 6 motion parameters was greater than 0.2 were censored. Volumes were also censored if more than 10% of the voxels in a volume contained large signal fluctuations that were considered outliers. Subject had 0-40 censored volumes, across all 4 pre and post TBS resting runs (1200 volumes total). Censored volumes per 300 volume run were: mean=2, median=0, maximum=22. The motion parameters and their first derivatives were regressed from the time series. In the same step, signals from a Cerebral Spinal Fluid (CSF) ROI and a white matter ROI that excluded voxels within 8 voxels of either stimulation site were averaged and regressed from the data. The resting-state data were bandpass filtered (0.01-0.1Hz) with censoring and the data were spatially smoothed with a 5-mm full-width-half-maximum Gaussian kernel.

### ROI specifications and cross-subject alignment

The statistical tests visualized in Figures 1 and 2 used regions of interest that were defined for each volunteer. The average of each volunteer’s anatomical scans, as described in the previous section, were processed in Freesurfer (Fischl et al., 2004) to estimate volunteer-specific anatomical boundaries for the fusiform gyri, lateral occipital gyri, and amygdala.

Any fMRI voxel that included at least 50% the Freesurfer estimated region was included in the region. These anatomical regions were intersected with faces>objects functional activation maps from the localizer scans collected during the TBS sessions (false-discovery-rate threshold q<0.05). To make sure ROIs for each brain region were identified in each volunteer, if 10% of the voxels in the anatomical ROI, or 10 voxels for anatomical ROIs with less than 100 voxels did not cross this threshold, then the threshold was increased until this threshold of faces>objects voxels was crossed. In 5 subjects, this liberal threshold was not crossed in the amygdala ROIs, which has a mean of 51 voxels in the anatomical ROIs. In 4 of these subjects, the same volunteers participated in a separate study that included the identical functional localizer task with more significant faces>objects voxels in the regions of interest (Pitcher et al., 2017). Those localizer scans were used to define the functional ROIs in those 4 volunteers. The voxels in the fusiform gyrus with the face>objects contrast were designated the Fusiform Face Area (FFA) ROI and the voxels in the lateral occipital gyrus were designated the Occipital Face Area (OFA) ROI. All volunteers had functionally localized rpSTS and rFFA ROIs with q<0.05. The threshold was q>0.05 for 1 volunteer in lFFA and rOFA, 2 volunteers in lpSTS, 3 volunteers in lOFA, 4 volunteers in rAmyg, and 8 volunteers in lAmyg. The left and right hand knob motor ROIs were hand drawn on each volunteer’s structural scan by a single person and checked for accuracy by a second person.

For whole-brain group maps, the within-subject aligned anatomical volumes from each subject were registered to a standard MNI space and then nonlinearly, iteratively warped to each other to make a within-study anatomical template (Figures 3 and S1).

## Author Contributions

D.A.H, P.A.B, L.G.U & D.P designed research; D.A.H, G.I, B.G, V.R, J.G-C & D.P performed research; D.A.H, G.C. & D.P analyzed data; D.A.H, P.A.B, L.G.U & D.P wrote the paper

## Acknowledgements

The research reported here was supported by the Intramural Research Program of the NIMH (ZIAMH002783 and ZIAMH002918) and by BBSRC grant BB/P006981/1 awarded to D.P. Clinical Protocol #: NCT01617408. This work utilized the computational resources of the NIH HPC Biowulf cluster (http://hpc.nih.gov). We thank Nancy Kanwisher for providing face localizer stimuli. We thank Shruti Japee for reviewing functional localizer results and for identifying functional localizer scans from these volunteers that were collected as part of other data sets.

