## Supplemental Figure 1 for "Thetaburst TMS to the posterior superior temporal sulcus decreases resting-state fMRI connectivity across the face processing network"

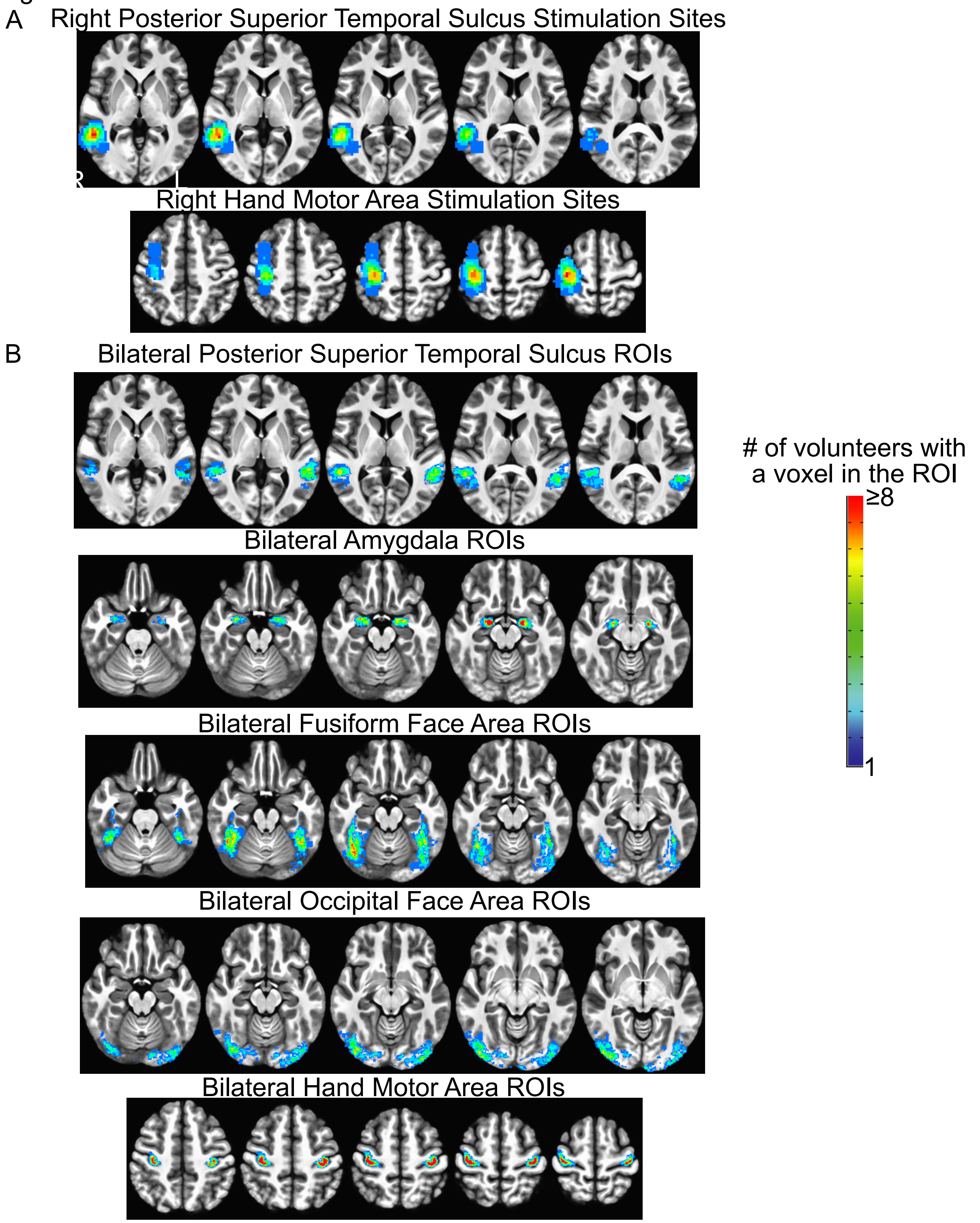


Union of the stimulation sites and ROI locations across volunteers. The anatomical underlay is the average of the aligned anatomical scans from the volunteers in this study. The overlay is the sum of all the volunter's ROIs. (A) The two stimulation sites across the subjects. The stimulation sites for each volunteer are visualized as a 1cm radius sphere centered on the stimulation site. The ROIs were individually localized for each stimulation site. The rpSTS stimulation site used a functional localizer and the motor stimulation site was defined using subject-specific anatomical landmarks. A re-examination of anatomical motor areas in each volunteer shows that the focus of the motor stimulation site was outside of the hand motor area for 5 volunteers with the focus being greater than 1 cm from the hand motor area in two volunteers. (B) Bilateral ROIs used for all connectivity analyses in this manuscript. All ROIs except for the hand motor areas were defined using the intersection of a Freesurfer-defined anatomical ROI and a Faces>Objects functional localizer
