## Supplemental Figure 2 for "Thetaburst TMS to the posterior superior temporal sulcus decreases resting-state fMRI connectivity across the face processing network"

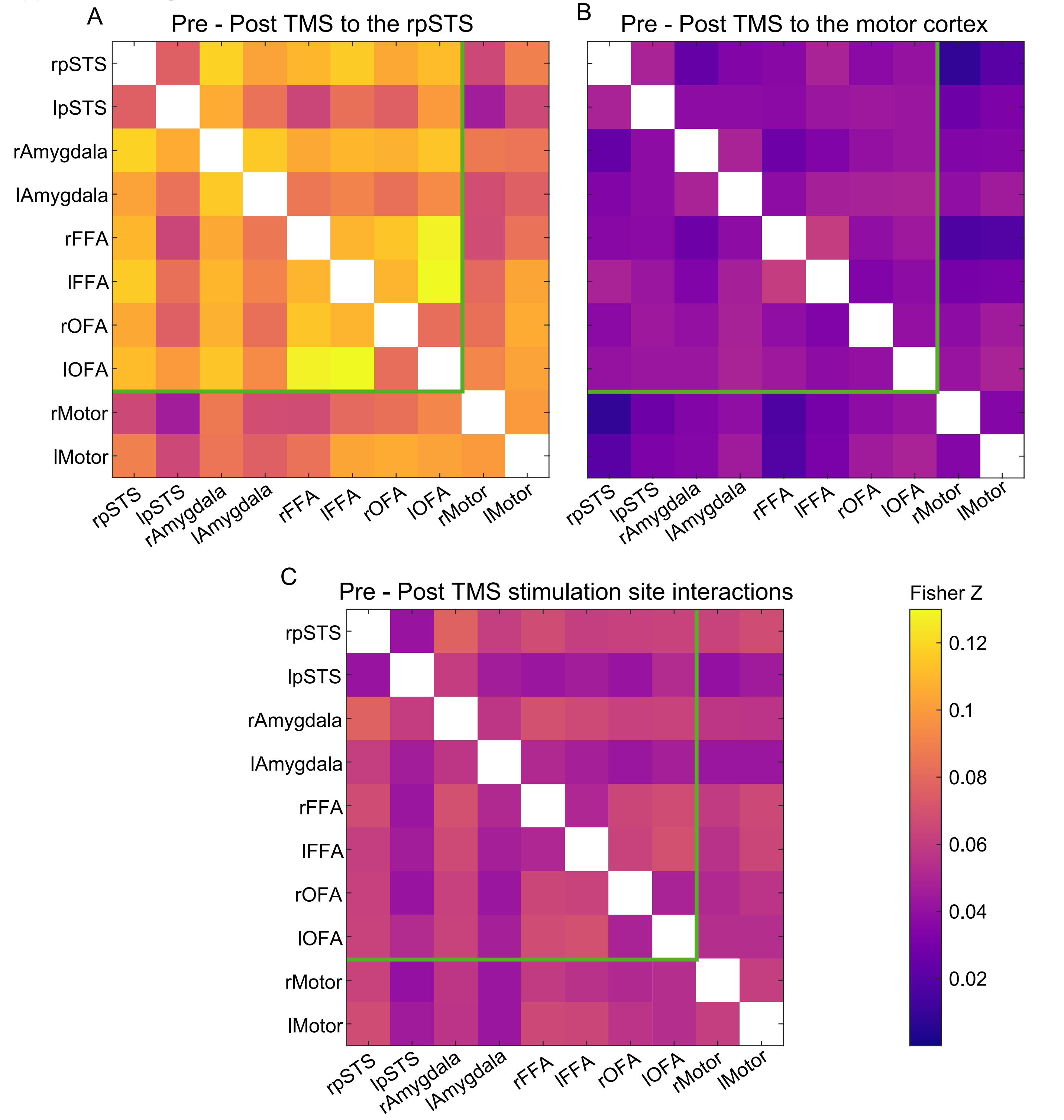


Fisher Z transformed correlation magnitude changes from pre - post TMS (A) To the rpSTS (B) To the the motor cortex, and (C) Pre - post stimulation & simulation site interaction effect. Magnitudes are the Matrix Based Analysis (MBA) model fits across the population. These are the results of the same analysis presented just for TMs to the rpSTS in Figure 1. Unlike Figure 1, the sizes of the squares are not scaled by their posterior probability. For the Motor cortex effect and the interaction effect, all posterior probabilities were less than 85%. The minimum value of the color scale is Z=0 rather than 0.2 in Figure 1 to better visualize the very small magnitude variations for motor stimulation and the interaction effects. The green line marks the ROI pairs that are within the pre-defined face-selective network.
