## Supplemental Figure 4 for "Thetaburst TMS to the posterior superior temporal sulcus decreases resting-state fMRI connectivity across the face processing network"

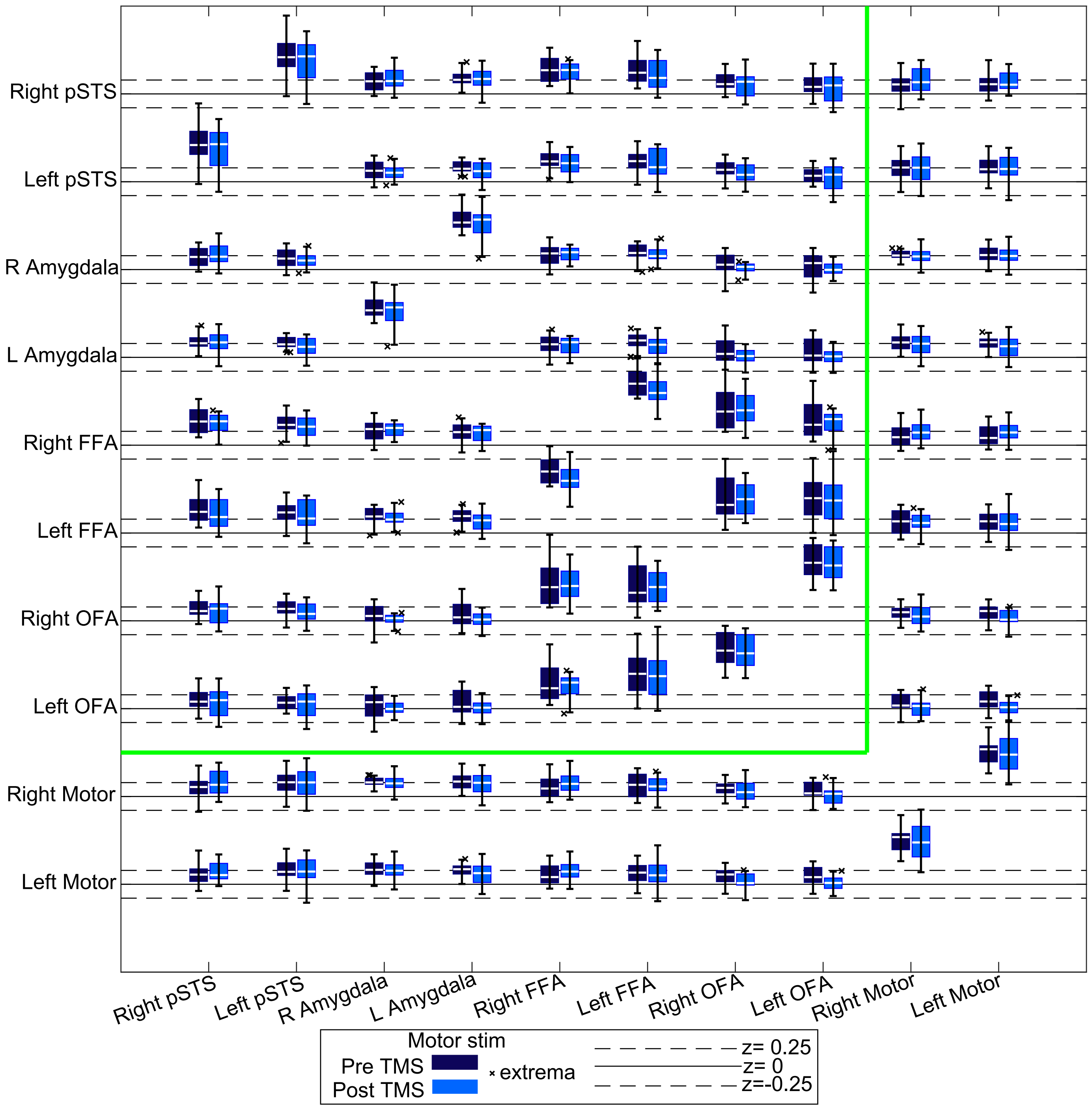


Correlations between the all pairs of ROIs in a pre-defined face network (within green line) as well as the bilateral primary motor hand regions. These boxplots correspond to the same data as the motor stimulation box plots in Figure 2 but for the entire matrix of ROI pairs as in Figure 1. The first row directly corresponds to the rpSTS correlations shown in Figure 2. The solid and dashed black lines demarcate -0.25<z<0.25 for each row. Magnitudes are the MBA model fits across the population for each condition. Boxplots show 25-75% of the distribution. The white line is the median. Whiskers are the maximum and minimum values excluding outliers. MBA was used to calculate posterior probabilites that a difference is greater than 0. None of the pre - post TMS to right motor cortex changes are above 85% posterior probability.
